# Integrated single-cell functional and molecular profiling of extracellular vesicle secretion in metastatic breast cancer

**DOI:** 10.1101/2022.08.13.503860

**Authors:** Mohsen Fathi, Melisa Martinez-Paniagua, Ali Rezvan, Melisa J. Montalvo, Vakul Mohanty, Ken Chen, Sendurai A Mani, Navin Varadarajan

## Abstract

Extracellular vesicles (EVs) regulate the tumor microenvironment by facilitating transport of biomolecular cargo including RNA, protein, and metabolites. The biological effects of EV-mediated transport have been studied using supra-physiological concentrations of EVs, but the cells that are responsible for EV secretion and the mechanisms that support EV secretion are not well characterized. We developed an integrated method based on arrays of nanowells to identify individual cells with differences in EV secretion and used an automated robot to perform linked single-cell RNA-sequencing on cloned single cells from the metastatic breast cancer cell line, MDAMB231. Gene expression profiles of clonal cells with differences in EV secretion were analyzed, and a four-gene signature of breast cancer EV secretion was identified: *HSP90AA1, HSPH1, EIF5*, and *DIAPH3*. We functionally validated this gene signature by testing it across different cell lines with different metastatic potential demonstrating that the signature correlated with levels of EV secretion. Analysis of the TCGA and METABRIC datasets showed that this signature is associated with poor survival, more invasive breast cancer types, and reduced CD8^+^ T cell infiltration in human tumors. We anticipate that our method for directly identifying the molecular determinants of EV secretion will have broad applications across cell types and diseases.

## Introduction

Extracellular vesicles (EVs) are a heterogeneous population of lipid bilayers that encapsulate and transport diverse biological cargo including nucleic acids and proteins^1,2^. EVs appear to be secreted from diverse mammalian cell types and are hypothesized to mediate long-range cell-cell communication^3,4,5^. In the last two decades, it has become evident that EV-mediated transport and delivery of biomolecules is important not only in normal physiological processes but also in pathological processes including cardiovascular disorders and cancers^6^. EVs influence every step of tumor progression and metastasis. Tumor-derived EVs induce upregulation of angiogenesis-related genes and enhance endothelial cell proliferation^7^, facilitate immunosuppression through the transfer of PD-L1^8^, enable invasion through the activity of the matrix metallopeptidase 2 (MMP2)^9^, and seed the pre-metastatic niche by downregulating expression of cadherin-17 in lung^10^. Tumor-derived EVs have potential for diagnostic purposes, and methods have been developed to dissect the heterogeneity of EVs down to the single particle-level^11,12,13^.

The pathways that regulate the secretion and packaging of EVs are not completely understood^14^. EVs are classified based on their mode of release as ectosomes or exosomes. Ectosomes, or shedding microvesicles, are released through the outward budding of the plasma membrane. Exosomes, by contrast, are synthesized by the inward budding of the endosomal membrane, which leads to formation of early endosomes. The maturation of early endosomes results in the formation of multivesicular bodies (MVB), and the fusion of the MVBs with the plasma membrane leads to secretion of exosomes. Proteins associated with MVB sorting include components of the ESCRT complex^15,16^; and ESCRT-independent molecules such as sphingolipid ceramide^17^ and tetraspanin CD63^18^. Proteins associated with the fusion of MVBs with the plasma membrane include the SNAREs^19,20^ and the RAB family members RAB27A, RAB27B, and RAB7^21,22^. Despite this progress, the map of proteins that participate in EV secretion remains incomplete. Most studies have focused on profiling the cargo of EVs, but this profiling largely reflects information being transferred rather than the molecules responsible for EV secretion^23,24^.

To directly link secretion of EVs with underlying molecular properties at single-cell resolution, we integrated single-cell EV profiling and cloning with single-cell RNA sequencing (scRNA-seq) to enable unbiased discovery of the genes that influence EV secretion. We utilized the well-validated metastatic breast cell line, MDAMB2321, which has been known to secrete EVs that enhance migration and invasion of cancer cells^25^. By performing scRNA-seq analysis on individual cells that either secrete high or low amounts of EVs, we discovered a four-gene signature, *HSP90AA1, HSPH1, EIF5*, and *DIAPH3*, correlated with EV secretion. We experimentally validated this core signature in breast cell lines in vitro and confirmed that HSP90 inhibitors negatively regulate EV secretion. Based on analysis of the Cancer Genome Atlas (TCGA), high expression of the EV signature is strongly correlated with poor survival and low CD8^+^ T cell infiltration in breast cancer patients.

## Results

### Establishing monoclonal cell lines with heterogenous EV secretion

To directly analyze EV secretion by clonal cells, we utilized nanowell arrays^26^. We mapped the heterogeneity in EV secretion within the metastatic triple-negative breast cancer cell line, MDAMB231, with the aid of two EV markers known to be expressed in these cells, CD63 and CD81 (**Figure 1A**). Single-cell profiling demonstrated that individual cells secrete very different amounts of EVs (**Figure 1B**)^27^. To determine whether or not EV secretion is a stably inheritable property, we retrieved individual EV secretor (labeled MDAMB231-S) and non-secretor cells (MDAMB231-NS) using an automated robot and expanded them to establish clonal populations. After limited expansion (<20 generations), we evaluated secretion rate by single cells from these clonal populations and confirmed that individual cells from the MDAMB231-S population secreted more EVs than MDAMB231-NS cells at all time points tested (**Figure 1C**). Tracking the kinetics of EV secretion showed that the majority (>87%) of both MDAMB231-S and MDAMB231-NS cells secreted EVs continuously over the 6 hour period monitored (**Figure 1D**). Since EV secretion is associated with increased migration in metastatic breast cancer cells^28^, we compared the migratory potentials of MDAMB231-S and MDAMB231-NS populations using a wound healing assay. The MDAMB231-S cells were significantly more migratory than were the MDAMB231-NS cells (**Figure 1E**). Taken together, these results showed that we can identify cells with differences in EV secretion and that the EV secretion property is maintained upon clonal expansion.

**Figure 1.**
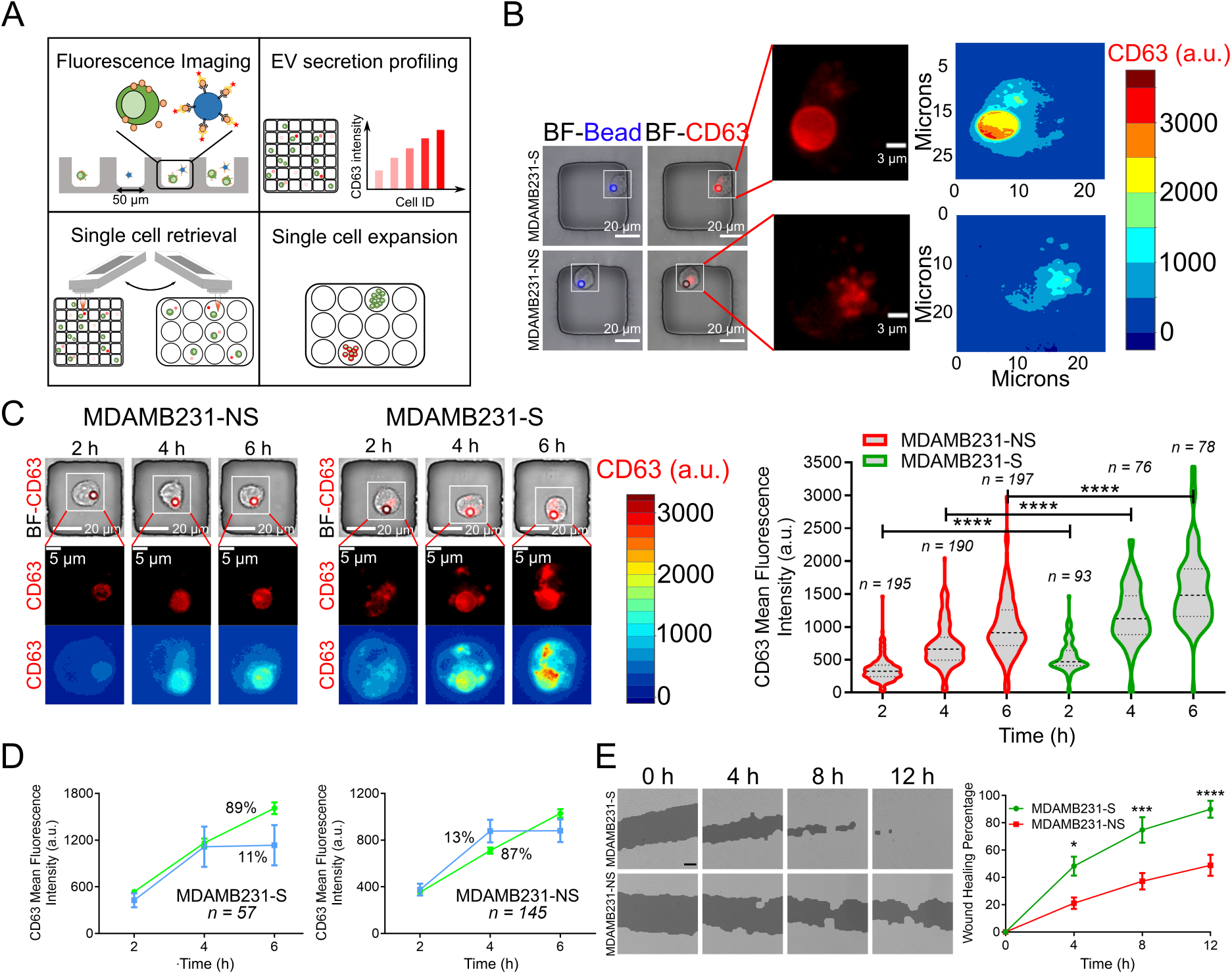
Establishment of monoclonal cell lines with different rates of EV secretion. A. The workflow for the identification and isolation of single cells with differences in EV secretion capacity. B. Representative images of MDAMB231-S and MDAMB231-NS cells in wells (left), at high resolution (middle), and as contour maps of intensity of CD63. C. Images of (top to bottom) cells in wells, high resolution of CD63, and CD63 intensity contour maps of representative MDAMB231-NS (left) and MDAMB231-S (middle) cells at 2, 4, and 6 h. Right: Violin plots of medians and quantiles of CD63 intensities (**** *p* < 0.00001; t-test). D. CD63 intensity as a function of time for MDAMB231-S (left) and MDAMB231-NS (right) cells plotted as means ± SEM. Two subpopulations are present: cells that continuously secrete EVs (green) and cells that secrete EVs in a burst at 2 h (blue). E. Left: Representative photographs of wound healing assays. Scale bar is 100 μm. Right: Plot of mean percent wound healed ± SEM versus time (n= 7 for each cell line; * *p* < 0.05, *** *p* < 0.001, and **** *p* < 0.0001; two-way ANOVA).

### Identification of an EV gene signature in breast cancer cells

The availability of the clonal populations allowed us to compare the transcriptional differences across thousands of single cells by scRNA-seq. To derive a genetic signature associated with EV secretion, we performed scRNA-seq on cells from the MDAMB231-S and MDAMB231-NS populations using the Rhapsody platform (**Figure 2A**). After data processing and filtering, we identified 1970 single cells with an average of 4710 unique genes and 24,219 transcripts per cell (**Figure S1A**). Dimensionality reduction showed a clear separation between the two cell types (**Figure S1B**); two clusters one consisting exclusively of secretor cells and one of non-secretor cells, were identified (**Figure 2B**). Differential gene expression analysis identified 322 genes were significantly enriched in the MDAMB231-S cells compared to MDAMB231-NS cells (adjusted *p* < 0.05; **Table S1**). We compared the differentially expressed genes (DEGs) to those previously associated with EV secretion^29,30,31^. Of the 322 DEGs we identified, 211 were annotated in the ExoCarta database as associated with EVs (**Figure 2C, Table S2**). When restricted to DEGs with greater than 1.2-fold difference between MDAMB231-S and MDAMB231-NS cells, we identified 34 genes known to be associated with EV secretion, metastasis and invasion (**Figure 2D, E, Table S3**). We classified the DEGs into genes that encode cell-adhesion/migration-related proteins (*e.g., ACTN1, CAV1, FXYD5*, and *DIAPH3*), transcriptional regulators (*e.g., GTF3A, TFDP1*,and *SNAPC1*), and chaperones and heat shock proteins (*e.g*., *HSP90AA1, HSPH1*, and *POMP*). Of these DEGs, *UBL3* encodes a protein that directly interacts with CD63 and functions as a key post-translational modifier that facilitates the sorting of proteins into EVs^32^.

**Figure 2.**
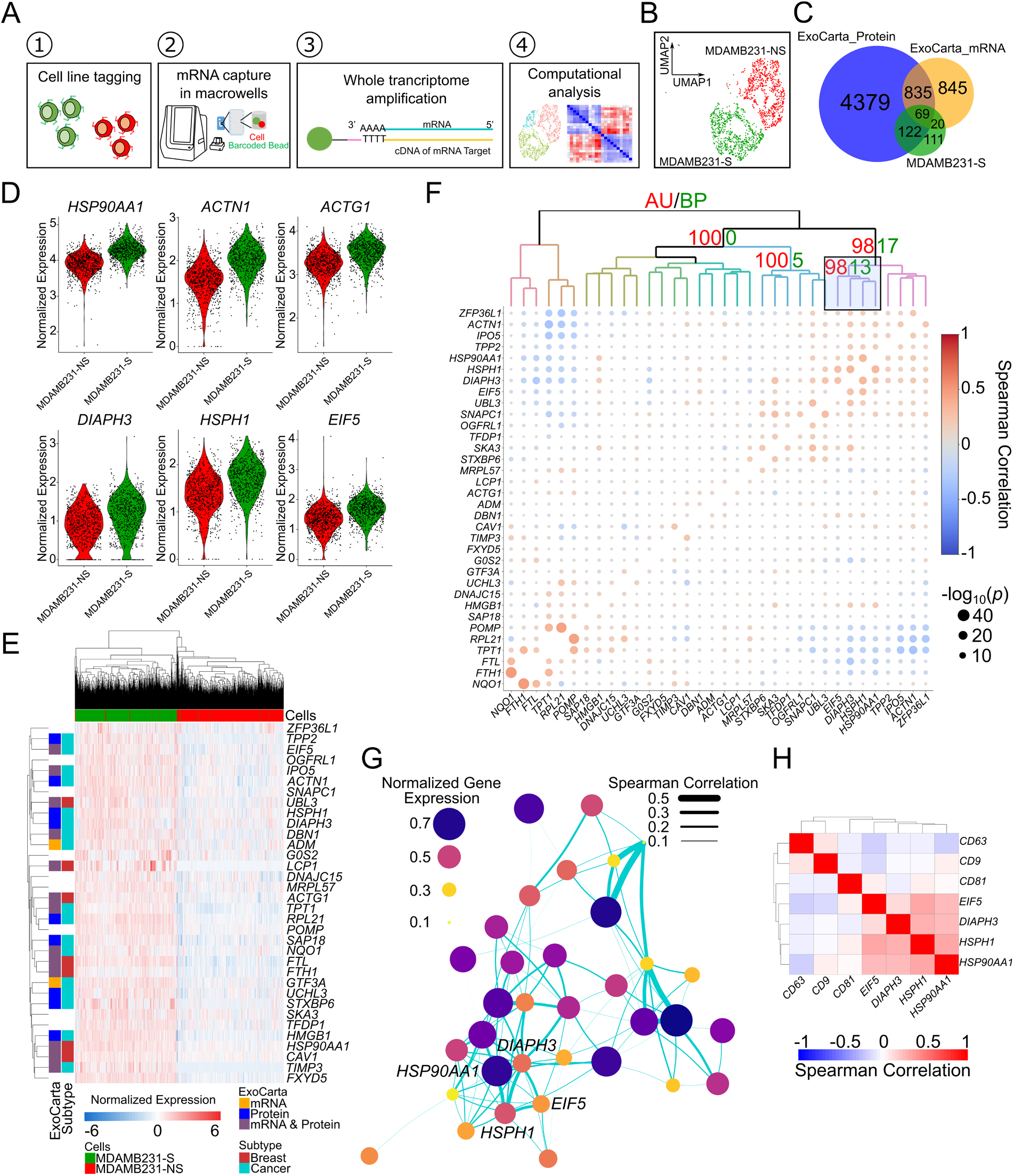
Identification of molecular signatures of EV secretion by scRNA-seq analysis. A. The workflow of single-cell RNA sequencing and whole transcriptome profiling for monoclonal cell lines. B. The UMAP plot of scRNA-seq data from MDAMB231-S and MDAMB231-NS cells. C. Venn diagram of the overlap of genes differentially expressed MDAMB231-S cells compared to MDAMB231-NS cells with mRNA and proteins from the ExoCarta dataset that are annotated as associated with EVs. D. Violin plots of genes upregulated in MDAMB231-S in comparison to MDAMB231-NS cells. E. Heatmap of the top 34 genes upregulated in MDAMB231-S cells. The colors to the left indicate ExoCarta annotation as associated with EV or linkage to breast cancer or other cancer types. F. Heatmap of Spearman coefficients for correlations between genes upregulated in MDAMB231-S cells relative to MDAMB231-NS cells. AU and BP indicate approximately unbiased and bootstrap probabilities, respectively. The correlations among EV-sig genes, *HSP90AA1, HSPH1, DIAPH3, and EIF5*, are highlighted. G. Network of top genes upregulated in MDAMB231-S cells relative to MDAMB231-NS cells with *HSP90AA1, HSPH1, DIAPH3, and EIF5* highlighted. H. Spearman correlation coefficients for four core EV genes and surface markers *CD63, CD81*, and *CD9* in MDAMB231-S cells.

We posited that the genes associated with EV secretion or packaging within the EVs are regulated in a coordinated manner. Accordingly, we calculated the Spearman coefficient between the DEGs and applied a hierarchical clustering to identify the cluster of genes that were significantly correlated with each other. By applying a multiscale bootstrap resampling method, we that *HSP90AA1* was significantly correlated with *HSPH1, EIF5*, and *DIAPH3* (**Figure 2F, G**). We refer to these four genes as EV-sig genes. The mRNAs encoded by each of these four genes were significantly correlated with the levels of EV marker *CD81* mRNA (**Figure 2H**). The proteins encoded by each of these genes have been individually shown to be associated with actin remodeling and EV secretion. *HSP90AA1* encodes HSP90, a molecular chaperone that promotes structural maintenance of proteins involved in cell cycle control and signal transduction. *HSPH1* encodes a member of the heat shock protein 70 family of proteins that acts as a nucleotide exchange factor for molecular chaperones and is capable of direct protein-protein interaction with HSP90^33^. *DIAPH3* encodes a member of the diaphanous subfamily. DIAPH3 is involved in actin remodeling and regulation of the cell motility and adhesion and can activate the beta-catenin/TCF signaling by binding to HSP90, which results in growth, migration, epithelial-mesenchymal transition, and metastasis in hepatocellular carcinoma cells^34^. *EIF5* encodes the eukaryotic translation initiation factor eIF5, which is enriched in EVs secreted from breast cancer cells and melanoma cells^35^.

To directly demonstrate a role for the EV-sig genes in EV secretion, we focused on HSP90 for three reasons: First, HSP90 interacts directly with both DIAPH3 and HSPH1 ^33,34^. Second, independent proteomic analyses of breast cancer cells showed that EVs contain both HSP90 and EIF5A^32^. Third, inhibitors of HSP90 are readily available, allowing us to study how inhibition of HSP90 impacts EV secretion from MDAMB231 cells. We used a standard transwell assay to capture the EVs secreted from MDAMB231 cells incubated with either tanespimycin (also known as 17AAG), a first generation HSP90 inhibitor, or ganetespib (also known as STA-9090), a potent, synthetic resorcinol-based HSP90 inhibitor (**Figure S2A**). The inhibition of HSP90 significantly reduced EV secretion in a dose-dependent manner even at nanomolar concentrations (**Figure S2B**). This observation further supports our hypothesis that EV-sig gene expression is a determinant of EV secretion.

### EV-sig is predictive of EV secretion in breast cancer cell lines

To validate whether EV-sig can predict the secretion of EVs, we investigated three breast cancer cell lines, MDAMB231, MCF7, and HCC70, which have differences in metastatic potential. MDAMB231 and HCC70 cell lines were established from triple-negative breast cancers, whereas the MCF7 line is an estrogen receptor- and progesterone receptor-positive cancer cell line^36^. The wound healing assay confirmed that the migration potential of MDAMB231 cells is significantly higher than those of MCF7 and HCC70 cells (**Figure 3A**). We performed scRNA-seq on MDAMB231, MCF7, and HCC70 cells. We obtained an average of 4459 unique genes and 22,071 transcripts per cell (**Figure S3A**). After dimensionality reduction, the cells from each of the three cell lines clustered separately (**Figure 3B**), and a total of 2634 DEGs (> 1.2-fold change) were identified (**Table S4**).

**Figure 3.**
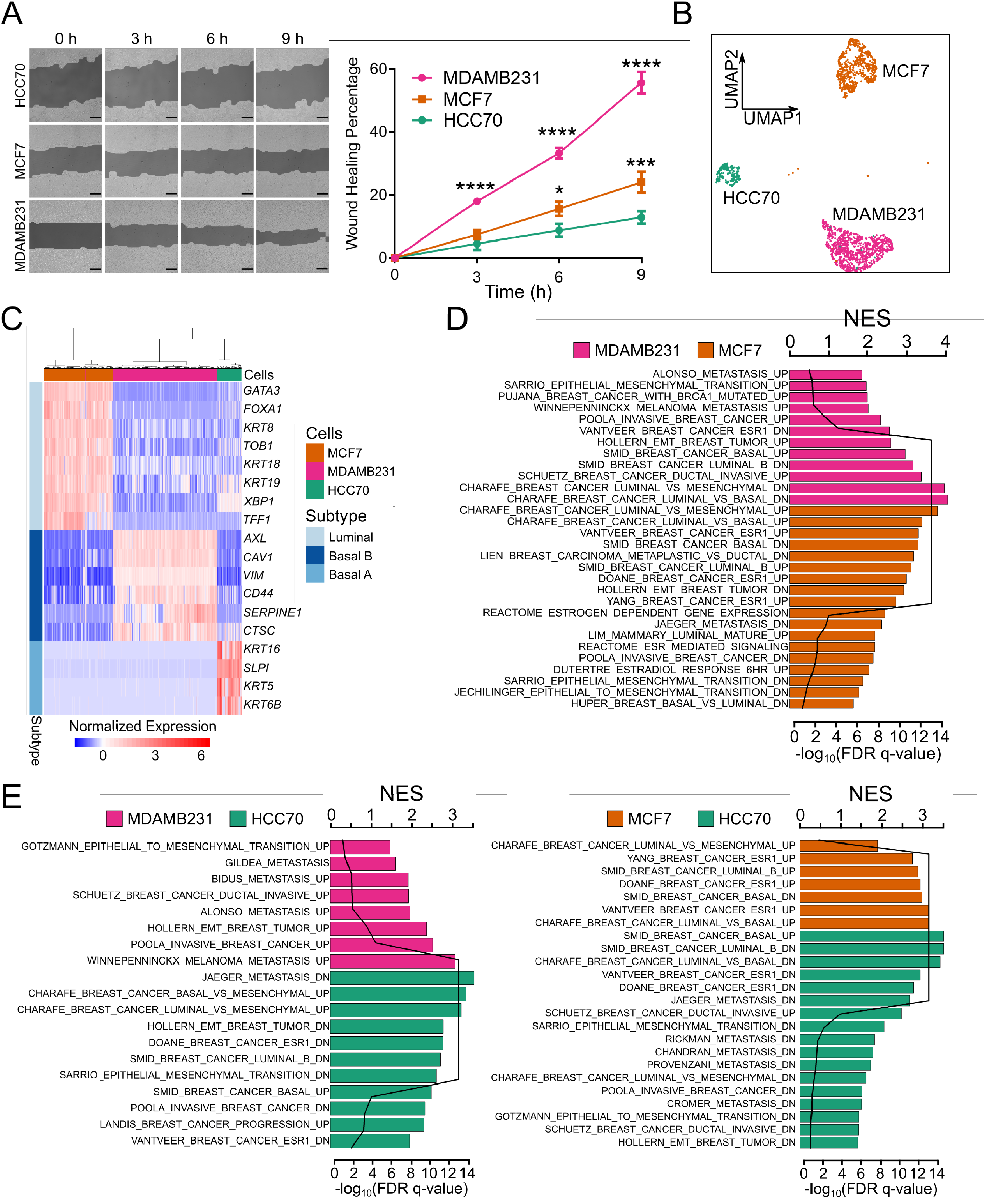
EV secretion is correlated with migration in breast cancer cell lines. A. Left: Representative images of wound healing assays showing the migration of MDAMB231, MCF7, and HCC70 cells over time. Scale bar is 100 μm. Right: Plot of mean percent wound healed ± SEM versus time (n=9; * *p* < 0.05, *** *p* < 0.001, and **** *p* < 0.0001; two-way ANOVA). B. UMAP plot of scRNA-seq data from MDAMB231, MCF7, and HCC70 cells. C. Heatmap of expression of genes associated with luminal, basal A and, basal B breast cancer subtypes in MDAMB231, MCF7, and HCC70 cell lines. D. Normalized enrichment scores (NESs) of pathways associated with metastatic cancer and luminal and basal breast cancer subtypes by pairwise comparison between MDAMB231 and MCF7 cells. E. NESs of pathways associated with metastatic cancer and luminal and basal breast cancer subtypes by pairwise comparisons between MDAMB231 and HCC70 cells (left) and between MCF7 and HCC70 cells (right).

To validate the phenotype of the cancer cell lines in the scRNA-seq data, we compared the expression of DEGs with known markers for breast cancer subtypes including luminal, basal A, and basal B^36^. This analysis showed that markers for luminal (e.g., *GATA3, FOXA1, KRT18*, and *KRT19*), basal A (e.g., *SLPI, KRT16*, and *KRT6B*), and basal B (e.g., *AXL, CAV1, VIM*, and *SEPRINE1*) subtype were upregulated in MCF7, HCC70, and MDAMB231 cells, respectively (**Figure 3C**). Similarly, pathway analysis confirmed that MDAMB231 and MCF7 cells are enriched for genes enriched in pathways corresponding to basal and luminal phenotypes, respectively (**Figure 3D**). Consistent with the fact that the HCC70 line was derived from a primary tumor, pathway analysis showed lower scores for metastatic and epithelial-mesenchymal transition pathways in HCC70 cells compared to the MDAMB231 and MCF7 cells (**Figure 3E**).

Next, we compared the average expression of EV-sig genes: HCC70 cells had the lowest expression, MCF7 cells had intermediate expression, and MDAMB231 cells had the highest expression (**Figure 4A**). Consistent with this observation, the Spearman correlation between the genes of the signature showed a significant correlation in the MDAMB231 cells; correlations were smaller in MCF7 and HCC70 cells (**Figure 4B**). At the protein level, we confirmed that all three cell lines expressed the HSP90 protein (**Figure S3B**). As an independent method to track the abundance of the EVs, we compared the levels of *CD63* and *CD81* mRNAs within the scRNA-seq data of these cell lines. All three lines expressed *CD63*; MDAMB231 and MCF7 cells expressed considerably higher levels of *CD81* mRNA than did HCC70 cells (**Figure 4C, S3C**).

**Figure 4.**
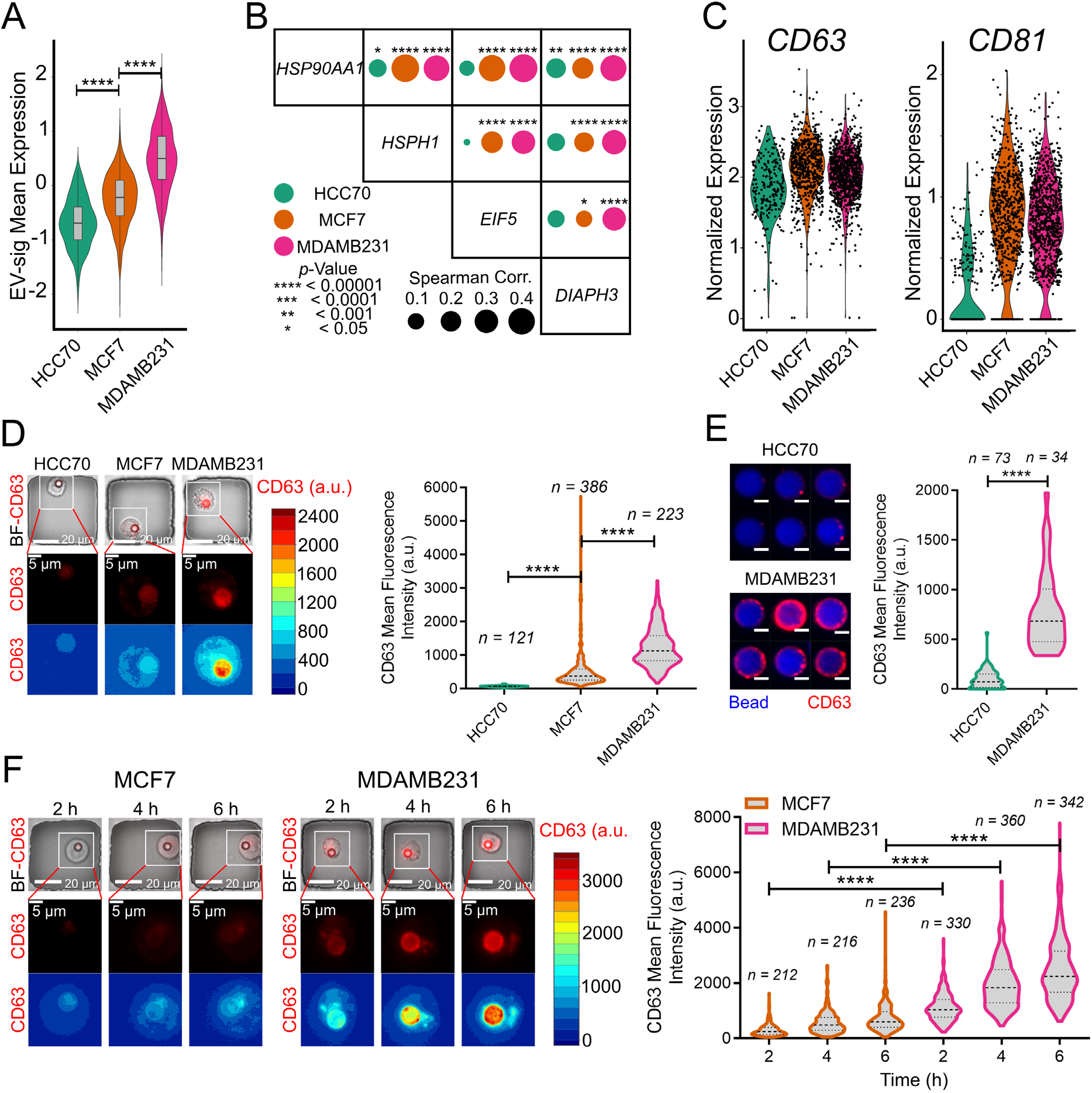
EV-sig is predictive of EV secretion by immortalized cells. A. Violin plots of average expression of EV-sig genes in MDAMB231, MCF7, and HCC70 cells (n = 227, 645, 971 for HCC70, MCF7, and HCC70 cells, respectively; **** *p* < 0.00001; t-test). B. Spearman correlation coefficients among EV-sig genes in MDAMB231, MCF7, and HCC70 cells. C. Violin plots showing the expression of *CD63* and *CD81* in MDAMB231, MCF7, and HCC70 cells. D. Left: Images of (top to bottom) cells in wells, high resolution of CD63 on cell, and CD63 intensity contour maps of representative MDAMB231, MCF7, and HCC70 cells at 6 h. Right: Violin plots of represent the median and quantiles of CD63 intensity (**** *p* < 0.00001, t-test). E. Left: Images of representative functionalized beads in culture with MDAMB231 and HCC70 cell lines for 48 h in transwell assay. Right: Violin plots of the median and quantiles of CD63 intensities (**** *p* < 0.0001; t-test). F. Images of (top to bottom) cells in wells, high resolution of CD63, and CD63 intensity contour maps of representative MCF7 cells (left) and MDAMB231 cells (Middle) at 2, 4, and 6 h. Right: Violin plots of the median and quantiles of CD63 intensities (**** *p* < 0.00001; t-test).

We then utilized our single-cell assay to directly profile EV secretion from each of these three cell lines. As predicted by EV-sig, HCC70 cells secreted low amounts of EVs, MCF7 cells secreted intermediate levels, and MDAMB231 cells secreted high levels (**Figure 4D**). We also independently validated these results using the standard transwell assay (**Figure S2A**), and these results confirmed that HCC70 cells secreted fewer EVs than did MDAMB231 cells (**Figure 4E, S4**). We tracked the short-term kinetics of EV secretion, and at all the timepoints tested, individual MDAMB231 cells showed higher EV secretion than did single MCF7 cells (**Figure 4F**). Lastly, we tested the impact of HSP90 inhibitors demonstrating that both tanespimycin and ganetespib inhibited EV secretion from MCF7 cells (**Figure S2**). In summary, scRNA-seq and EV profiling results showed that more migratory cells secrete more EVs than do less migratory cells and that EV-sig can predict cells amount of EV secretion.

To generalize the value of EV-sig, we obtained gene expression data on 1304 cell lines available in the Broad Institute Cancer Cell Line Encyclopedia (CCLE). Within this expanded dataset, expression of the EV-sig genes was highly correlated (**Figure S5A**). Focusing specifically on breast cancer, EV-sig showed highest expression in basal B phenotypes, followed by basal A, then HER2-enriched, and then luminal (**Figure S5B**). This is consistent with the known aggressiveness of these subtypes of breast cancer.

### EV-sig correlates with breast cancer outcomes

To investigate the translational value of EV-sig, we took advantage of the breast cancer datasets available in TCGA and METABRIC. We interrogated combined transcriptomic and clinical/pathological annotations for 1093 and 809 patients with breast cancer in TCGA and METABRIC, respectively. We first confirmed that the four genes that comprise EV-sig are significantly correlated with each other within human breast cancers (**Figure 5A, S6A**). We then stratified the patients into two groups: those with high EV-sig expression (BRCA_EV^Hi^ and MET_EV^Hi^) and those with low EV-sig expression (BRCA_EV^Lo^ and MET_EV^Lo^). The overall survival of patients with EV^Hi^ tumors was significantly lower than patients with EV^Lo^ tumors. For patients with data in TCGA, the median survival for patients with EV^Hi^ was 7.5 years, whereas those with EV^Lo^ the median survival was 10.8 years (HR: 2.3, 95% CI: 1.53-3.45); for patients with data in METABRIC, the median survival was 49.4 months for patients with EV^Hi^ and 63.6 months for EV^Lo^ (HR: 1.2, 95% CI: 1.03-1.40) (**Figure 5B, S6B**). Quantification of the pathology of the disease showed that EV-sig was associated with increased tumor size and more advanced disease (**Figure 5C-D, S6C**).

**Figure 5.**
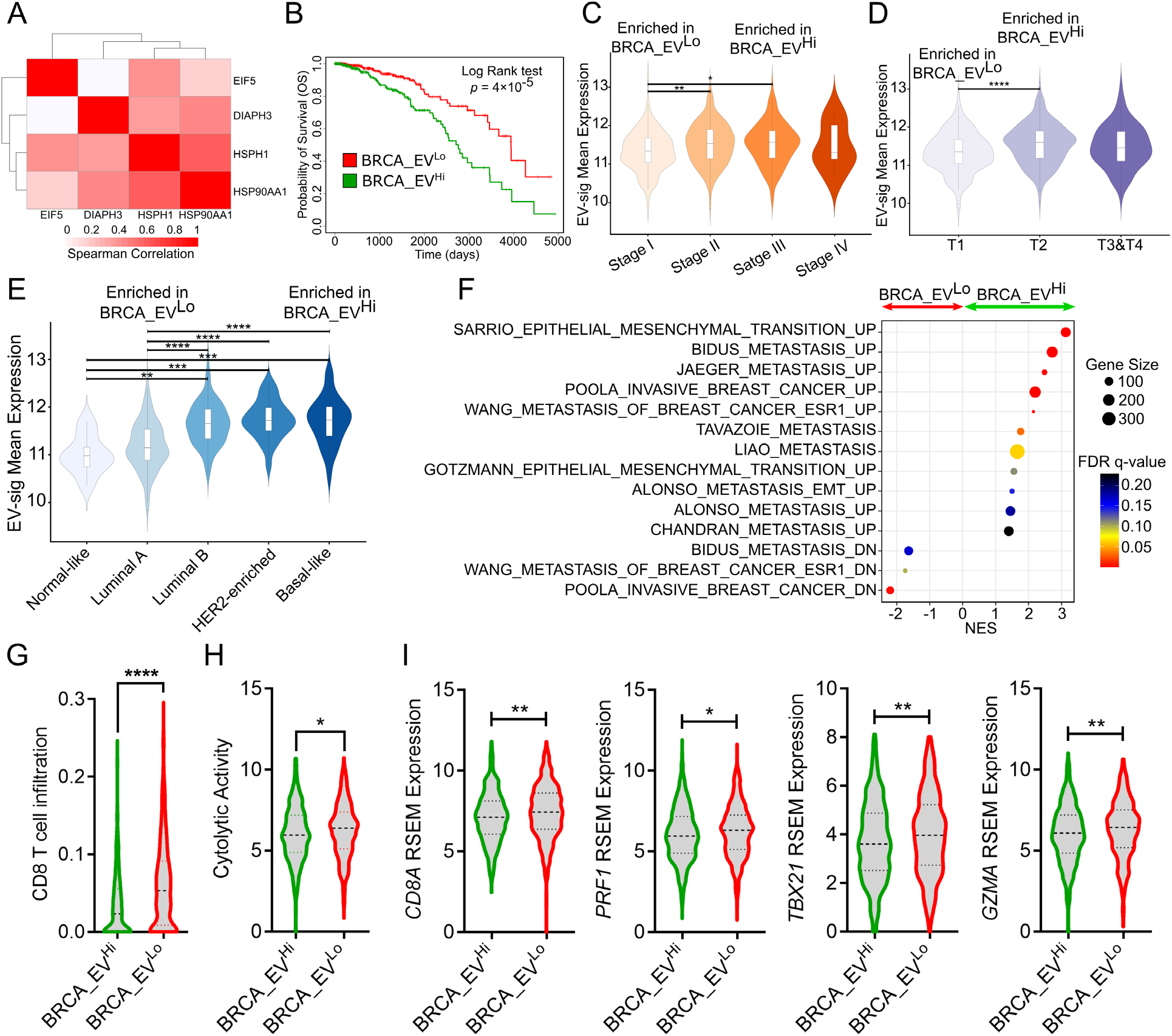
Expression of EV-sig genes is correlated with poor survival in breast cancer patients. A. Spearman correlation coefficients of EV-sig genes in breast cancer patients from TCGA dataset. B. Overall survival of breast cancer patients divided by the median of the average expression of EV-sig genes (n = 547, and 546 for BRCA_EV^Hi^ and BRCA_EV^Lo^, respectively). C. Violin plots of average expression of EV-sig genes by breast cancer stage (n=108, 371, 153, and 13 for Stage I to V, respectively; * *p* < 0.05, ** *p* < 0.01, *** *p* < 0.001, and **** *p* < 0.0001; one-way ANOVA). D. Violin plots of average expression of EV-sig genes by the size of the tumors with T1 the smallest (n= 174), T2 intermediate (n =395), and T3 & T4 the largest (n=94); (* *p* < 0.05, ** *p* < 0.01, *** *p* < 0.001, and **** *p* < 0.0001; one-way ANOVA). E. Violin plots of average expression of EV-sig genes by the breast cancer subtype (n=8, 191, 109, 53, and 83 for normal-like, luminal A, B, HER2-enriched, and basal-like subtypes, respectively; * *p* < 0.05, ** *p* < 0.01, *** *p* < 0.001, and **** *p* < 0.0001; one-way ANOVA). F. NESs of pathways associated with metastasis in patients with high and low levels of EV-sig gene expression. G. CD8^+^ T cells infiltration based on CIBERSORTx data for patients with high and low levels of EV-sig gene expression (**** *p* < 0.0001; t-tests). H. Cytolytic activity scores and geometric means of *PRF1* and *GZMA* mRNA levels in patients with high and low levels of EV-sig gene expression (* *p* < 0.05; t-test). I. Normalized expression of CD8^+^ T cell gene signature (*CD8A, PRF1, TBX21, GZMA*) in the patients with high and low levels of EV-sig gene expression (* *p* < 0.05, ** *p* < 0.01, *** *p* < 0.001, and **** *p* < 0.0001; t-test).

We next evaluated whether EV-sig could be used to stratify the different molecular subtypes of breast cancers. The tumors with low levels of EV-sig expression were enriched in normal-like and luminal breast cancers, whereas tumors with high levels of EV-sig expression were enriched in basal breast cancers (**Figure 5E, S6D**). Gene set enrichment analysis (GSEA) specifically focused on pathways associated with tumor cell functions showed that tumors with high levels of EV-sig expression are enriched in pathways associated with invasiveness, metastasis, and epithelial to mesenchymal transition (**Figure 5F, S6E**). Taken together, the clinicopathological data are consistent with our in vitro observation that EV secretion is associated with increased aggressiveness and invasion of tumor cells.

To evaluate the nature and frequency of immune cell infiltration associated with higher expression of EV-sig genes, we quantified the relative frequencies of the 22 different immune cell types by evaluation of normalized gene expression data using the CIBERSORTx algorithm. Tumors classified as EV^Lo^ had an increased frequency of CD8^+^ T cells, increased cytolytic activity (associated with increased expression of both *PRF1* and *GZMA*), and increased frequency of *TBX21* expression in comparison to EV^Hi^ tumors (**Figure 5G-I, S6F**). Collectively, these results suggest that increased EV secretion by cancer cells is associated with decreased CD8^+^ T cell infiltration and that this in turn promotes growth of larger and more aggressive tumors.

## Discussion

EVs secreted by cancer cells have potential to serve as diagnostic markers, and EVs could be used for delivery of anticancer agents into tumors^37,38,39^. Techniques based on analysis of bulk cells have enabled classification of EVs, characterization of the cargoes of EVs, and investigation of the impact of EVs on the progression of tumors^40^. At the other end of the spectrum, single-vesicle profiling studies have revealed the heterogeneity of both surface markers and the internal cargo of EVs^41,42,43,44^. In the past few years, single-cell methods to map the heterogeneity in EV secretion across single cells have been developed^45,46,47^. We recently reported a method to profile secretion of EVs from single cancer cells using nanowell arrays^26^ and here used this methodology to perform integrated profiling of the EV secretion and the transcriptional signature of EV-secreting cancer cells.

By profiling of populations of metastatic MDAMB231 breast cancer cells with differences in EV secretion, we identified several transcripts that are differentially expressed in cells that do and do not secrete EVs. Several were previously shown to play key roles in the biogenesis and secretion of EVs. *CAV1* encodes a protein that blocks the fusion of MVBs with autophagosomes^48^. EVs containing CAV1 have been shown to enhance the proliferation and invasion of metastatic cells^49^. *UBL3* encodes a protein that was shown to enhance the sorting of cargo into EVs by functioning as a post-translational modification factor^32^. The majority of the proteins encoded by our DEGs were shown to interact UBL3 in MDAMB231 cells by unbiased proteomics, suggesting the importance of UBL3-mediated sorting of protein cargo into EVs^32^. Although initial studies suggested that the C-terminal CAAX motif, which is CVIL in UBL3, is important for membrane localization and substrate modification, how UBL3 identifies substrate proteins and mediates the transport of these proteins into EVs are not known.

Based on EV secretion analysis and scRNA-seq data from MDAMB231 cells, we derived a transcriptional signature of EV secretion, based on four genes, *HSP90AA1, HSPH1, EIF5*, and *DIAPH3*. The levels of these genes are strongly correlated in cancer cell lines. We validated this signature by utilizing EV-sig to predict the EV secretion propensities of breast cancer cell lines and by inhibiting the activity of HSP90 in vitro and confirming that inhibition of HSP90 activity reduces EV secretion. We further demonstrated that expression of EV-sig genes is associated with aggressiveness of 1304 cell lines available in CCLE. The abundance of HSP90 in cancer EVs has been extensively documented, and it was recently shown that HSP90 is expressed in the EVs of more than 80% of cancer cell lines^50^. HSP90 is a pivotal regulator of proteostasis in cancer cells due to the high stress burden of these cells. In addition to its role as a chaperone, HSP90 also mediates the fusion of MVBs with the plasma membrane in yeast cells directly leading to secretion of EVs^51^. Somewhat surprisingly, tanespimycin, which traps HSP90 in the open conformation, does not alter the membrane deformation (and presumably EV secretion) activity of *Drosophila* HSP90 expressed in yeast cells^51^. By contrast, our results directly evaluating EV secretion demonstrated that treatment of MDAMB231 cells with tanespimycin reduced EV secretion. The differences in these two results may be due to the differences in the readouts used to profile EV secretion: membrane deformation in the *Drosophila* cells vs. direct profiling of EV secretion here. The pivotal role for HSP90 in influencing secretion of EVs containing CD63 was independently confirmed via knockout of *HSP90^52^*.

To determine whether our EV-sig genes can act as biomarkers of EV secretion in patient biopsies, we investigated the correlation of EV-sig gene expression with outcome for patients whose data are available through The Cancer Genome Atlas (TCGA) and METABRIC databases. We discovered that elevated expression of EV-sig genes is associated with increased tumor size and stage of cancer and an enrichment of more aggressive subtypes such as basal-like and HER2-enriched subtypes. Not surprisingly, elevated expression of EV-sig genes was associated with poor survival, and immune decomposition analyses revealed that tumors that express high levels of these genes were characterized by poor CD8^+^ T cell infiltration.

In summary, we have performed integrated single-cell profiling of EV secretion and transcriptomes of breast cancer cells. We provided direct evidence for the role of HSP90 in EV secretion and showed that expression levels of four genes, which are correlated with EV secretion in cell lines, can be used to stratify tumor aggressiveness and survival within breast cancer patients. We anticipate that our method for directly identifying the molecular determinants of EV secretion will have broad applications across cell types and diseases.

## Material and Methods

### Cell culture

MDAMB231, HCC70, and MCF7 cells were purchased from ATCC. We cultured MDAMB231 and HCC70 cells in RPMI 1640 supplemented with 10% FBS, 1% L-glutamine, 1 % HEPES, and penicillin-streptomycin. We cultured MCF7 cells in Eagle’s Minimum Essential Medium with 10% FBS, 1% HEPES, MEM Non-Essential Amino Acids, and penicillin-streptomycin. We tested all cells for mycoplasma contamination using real-time PCR.

### Single-cell EV detection assay

We analyze the secretion of EVs from single cells as previously described^26^. Briefly, we labeled cells with PKH67 dye (Sigma-Aldrich, catalog number PKH67GL-1KT) as directed by the manufacturer. To capture the EVs on the surface of LumAvidin beads (Luminex, catalog number L100-L115-01), we incubated the beads with 3.5 μg/ml biotinylated anti-CD81 antibody (BioLegend, clone TAPA-1) at the room temperature for 40 min, followed by three washes in PBS with 1% BSA. Then we loaded the labeled cells and functionalized beads into nanowells coated with PLL-g-PEG (SuSoS) and incubated at 37 °C. At 45 min before each detection time point, we added 4 μg/ml PE-labeled anti-CD63 antibody (BioLegend, clone H5C6). We imaged the nanowell using Zeiss Axio Observer Z1 microscope equipped with 20x/0.8 NA objectives and a Hamamatsu Orca Flash v2 camera.

We analyzed the TIFF images exported from the microscope as previously described^26^. Briefly, we segmented, quantified the cell to bead ratio, identified the cell to bead ratio of 1:1, and calculated the background-subtracted pixel values for identification of secreting and non-secreting cells. To analyze the dynamic of secretion from single cells, we detected the wells which maintained the 1:1 ratio during entire time course of the experiment. Then, based on a two-tailed t-test of the CD63 intensity per pixel, we selected a significant increase with *p*-value < 0.01 as the criterion for a change in the secretion behavior of the cell.

### Establishment of clonal cell lines

As previously described^26^, we used the CellCelector micromanipulator (ALS) equipped with 50-μm glass capillaries to retrieve the detected secretor and non-secretor single cells. We transferred the retrieved cells to a 96-well plate containing complete media and cultured the cells up to 24 population doublings.

### Wound healing assay

We cultured MDAMB231-S and MDAMB231-NS cells to 90% confluency in 12-well plates in media containing 10% FBS. We then replaced the media with media containing 0.5% FBS and cultured for an additional 18 h. After this starvation period, we scratched the monolayer with 10 μl pipette tips and washed the wells twice with PBS. During the assay, we incubated the cells with media containing 0.5% FBS. At several time points, we obtained images with the Zeiss Axio Observer Z1 microscope equipped with 20x/0.5 NA objectives. We analyzed the images with the Tscratch tool^53^.

### EV quantification using a transwell assay

We utilized a Transwell insert with a 3-μm pore membrane. and loaded functionalized beads in the lower compartment, and cells in the upper compartment of the insert. For HSP90 inhibitor assays, we covered the transwell with 10 nM or 30 nM of tanespimycin or ganetespib. After 48 h at 37 °C, we collected the beads and labeled them with 4 μg/ml PE-labeled anti-CD63 antibody (BioLegend, clone H5C6) for 45 min at 37 °C. We subsequently washed the beads three times in PBS with 1% BSA and imaged the wells using a Zeiss Axio Observer Z1 microscope equipped with 20x/0.8 NA objectives. Using ImageJ, we segmented and measured the fluorescent intensity of CD63.

### Surface marker staining

To measure the expression of CD81, we coated the cells with 3.5 μg/ml biotinylated anti-CD81 (BioLegend, clone TAPA-1) antibody at the 37 °C for 30 min. After one wash in PBS with 1% BSA, we stained the cells with 4 μg/ml PE-streptavidin (BioLegend) at the 37 °C for 45 min. We imaged the cells using a Zeiss Axio Observer Z1 microscope equipped with 20x/0.8 NA objectives, and quantified the CD81 signal using ImageJ.

### Single-cell RNA-sequencing

We labeled HCC70, MCF7, MDAMB231, MDAMB231-S, and MDAMB231-NS cells separately with the Sample-Tags from the BD Human Immune Single-Cell Multiplexing Kit (BD Biosciences) as described in the manufacturer’s protocol. Then, we prepared a library from ~5000 cells (approximately 1000 cells from each group). We used the BD Rhapsody System to prepare samples for transcriptome analysis. We assessed the quality and quantity of the final library using the Agilent 4200 TapeStation system using the Agilent High Sensitivity D5000 ScreenTape and a Qubit dsDNA HS Assay, respectively. We diluted the final library to 3 nM concentration and used a HiSeq PE150 sequencer (Illumina) to perform the sequencing.

### Sequencing read alignments

We analyzed the FASTQ files using the BD Rhapsody WTA Analysis Pipeline available on the Seven Bridges website (https://www.sevenbridges.com/). After performing alignment, filtering, and sample tag detection, we downloaded the sample tag calls and molecule count information for further analysis in R (v 4.0.1) using Seurat Package (v 3.0)^54^.

### Data processing and identification of differentially expressed genes

We performed the clustering using the standard processing workflow in the Seurat Package. Briefly, we removed cells with less than 8000 gene count, cells with mitochondrial genes at greater than 20% of the reads, and cells in clusters that contained a mixture of sample tags, resulting in 3431 single-cell profiles (773 MDAMB231-S cells, 815 MDAMB231-NS cells, 971 MDAMB231 cells, 645 MCF7 cells, and 227 HCC70 cells). Next, we identified the differentially expressed genes using the *Findmarkers* function in Seurat. We selected the markers with greater than 1.2-fold higher expression in MDAMB231-S cells in comparison to MDAMB231-NS cells as the gene signature for EV secretion.

### ExoCarta dataset analysis

We downloaded the list of proteins and mRNAs in the ExoCarta dataset http://exocarta.org/download.

### Gene correlation analysis

To calculate the Spearman correlation between genes, we used the *cor.test* function in R. We created the heatmaps of correlation coefficients with *pheatmap* package (v 1.0.12).

### Gene set enrichment analysis for breast cancer cell lines

To perform pathway analysis, we pre-ranked DEGs (*p*-value < 0.05) between each pair of cell lines identified using the *Findmarkers* function in *Seurat* package. We ran the GSEA software provided by UC San Diego and Broad Institute using Broad Institute C2: curated gene sets.

### Core signature identification and network analysis

We calculated the Spearman correlations between identified DEGs between MDAMB231-S cells and MDAMB231-NS cells. We used *ward.D2* as a hierarchal clustering method along with *Euclidean* distance method to cluster the markers. Using the *pvclust* package (v 2.2-0)^55^, we assessed the uncertainty in clustering analysis. We used the approximately unbiased value > 95 as the criteria for a significant cluster. We plotted the heatmap using *pheatmap* package.

To build the network between markers, we used *igraph* package (v 1.2.5)^56^. First, we created an undirected network containing a list of links and nodes. The size of nodes represented the average gene expression of each marker and the links represented the Spearman coefficient between each marker. Next, we removed the negative links, and to simplify the network, we removed the links that showed a smaller coefficient than the average of positive links. To visualize the network, we used the layout algorithm of layout_with_graphopt.

### Cancer Cell Line Encyclopedia (CCLE) analysis

We downloaded the CCLE log2 transformed RNA-seq TPM gene expression and the cell line information from the DepMap portal (https://depmap.org/portal/download/). To perform the correlation analysis, we filtered the gene expression matrix for the genes of EV-sig and determined Spearman correlations among these genes in the 1304 cell lines available. To analyze the correlations with respect to breast cancer subtype, we first selected breast cancer cell lines using the *primary_disease* information. Then, using the *lineage_molecular_subtype*, we grouped the cell lines into different subtypes.

### TCGA and METABRIC analyses

We downloaded all the TCGA data, including raw counts, RSEM gene normalized expression, and clinical data from the Broad Institute FireBrowse Data Portal (www.firebrowse.org). For tumor size, and stage analyses, we downloaded the *BRCA_clinicalMatrix* file from University of California Santa Cruz Xena Hub Portal (https://xena.ucsc.edu/) and used *PAM50_mRNA_nature2012, Tumor_nature2012*, and *AJCC_Stage_nature2012* for PAM50, tumor size, and stages information, respectively. We downloaded all METABRIC data, including gene expression and clinical data from the cbioportal for cancer genome (www.cbioportal.org). We calculated the Spearman’s rank correlation coefficients using the *cor.test* function and plotted using *pheatmap* package in R. For survival analysis, we used the Kaplan-Meier method. We compared the overall survival of patients divided by the median expression of the four EV-sig genes. Using the log-rank test, we calculated the statistical significance of survival curves. To perform pathway analysis, we identified DEGs between patients and divided by the median expression of the four EV-sig genes using the *DESeq2* (v 1.22.2) package^57^ for TCGA dataset. To calculate the DEGs for the METABRIC dataset, we performed a Wilcoxon test. We next used the pre-ranked genes with a significant *p*-value of < 0.05 to run GSEA software provided by UC San Diego and Broad Institute using Broad Institute C2: curated gene sets. We used the normalized gene expression of breast cancer patients to estimate the relative fraction of 22 immune cell types using 1000 permutations with the CIBERSORTx analytical tool. We calculated the cytolytic activity as the geometric mean of *PRF1* and *GZMA* as previously described^58^.

## Author Contributions

N.V. and M.F. designed the study. M.F. performed *in vitro* study. M.M.P. and A.R. performed scRNA-seq. M.F. performed scRNA-seq analysis. M.F. and V.K. performed TCGA and METABRIC analysis. M.J.M. performed western blotting. K.C., S.A.M, and N.V. supervised the study.

## Supplemental Materials

Additional figures and tables (PDF)

Supplementary Tables; S1, S2, S3, and S4

