## Supplementary Materials for "Integrated single-cell functional and molecular profiling of extracellular vesicle secretion in metastatic breast cancer"

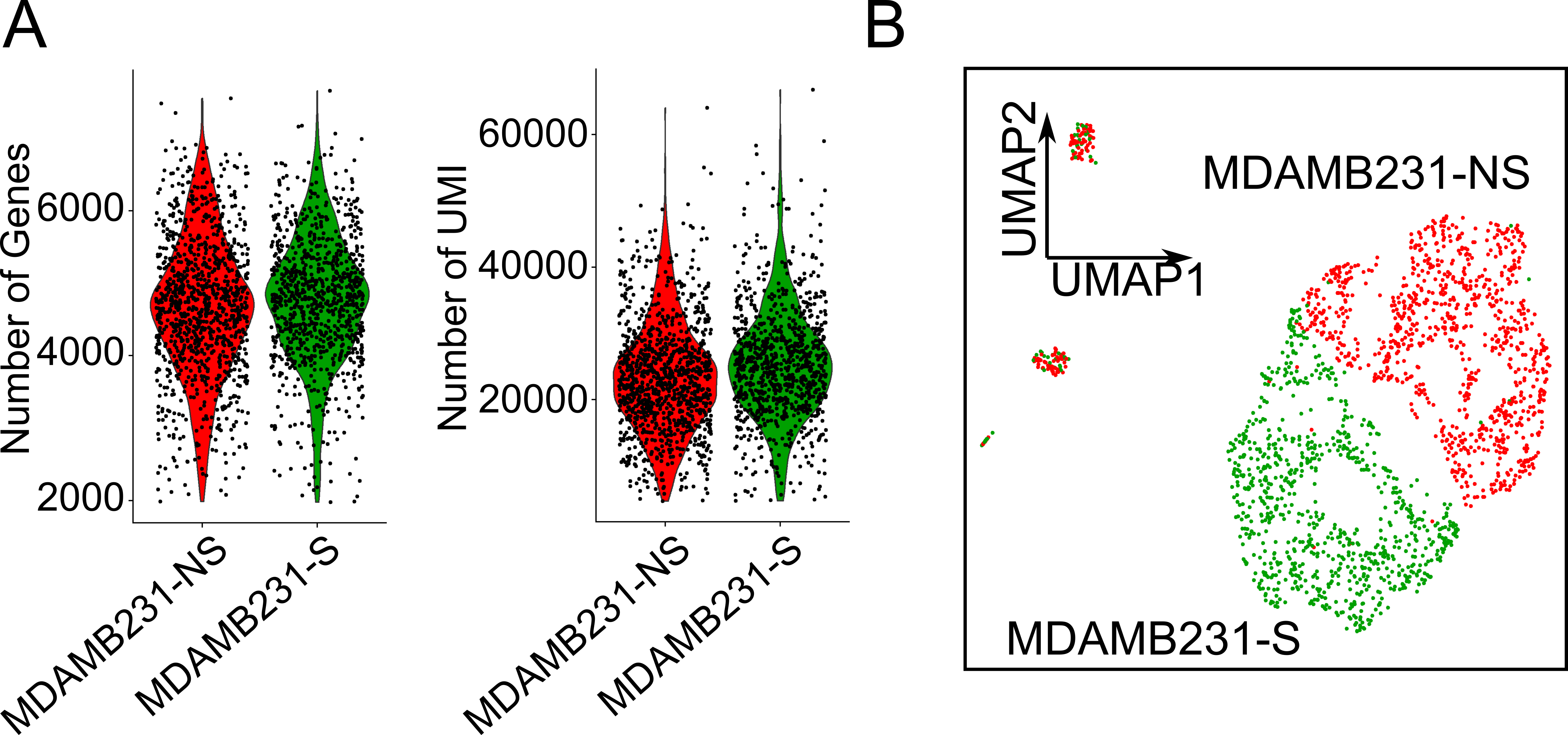

Figure S1. The scRNA-seq analysis of MDAMB231-S and MDAMb231-NS cell lines

A. Violin plot of the number of genes and transcriptome per cells.
B. UMAP plot of the MDAMB231-S and MDAMB231-NS cell lines.

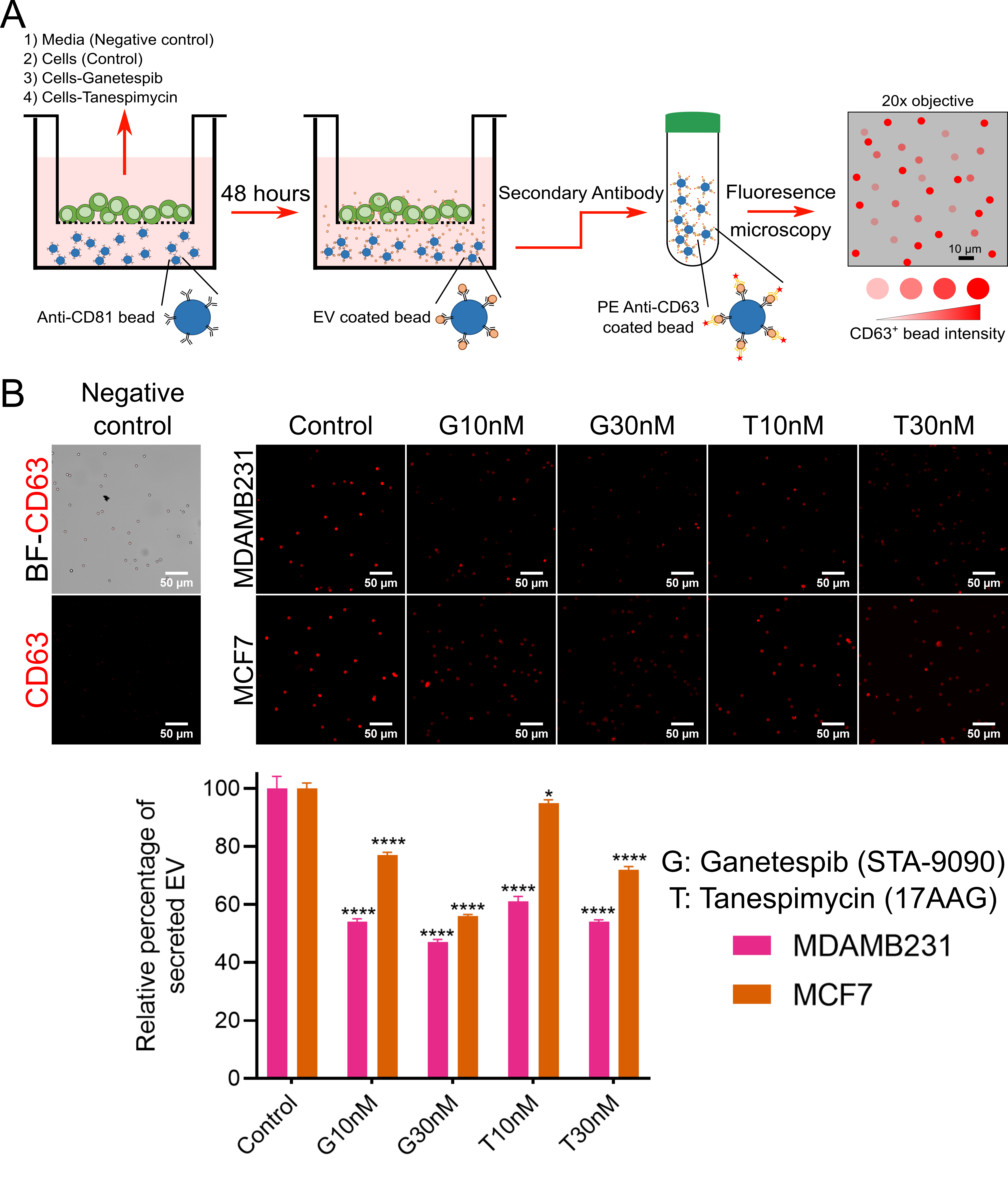

Figure S2. Decrease of EV secretion from MDAMB231 and MCF7 cells upon inhibition of HSP90

A. Workflow of transwell assay for quantifying the secretion of EV from population of cells.
B. Top: Fluorescence images of beads incubated with cells on the lower compartment of transwell inserts in the culture with or without HSP90 inhibitor (tanespimycin, and ganetespib) at 48 h. Bottom: Barplot of the relative percentage of secreted EV compared to non-treated cells (mean ± SEM) (* *p* < 0.05, and **** *p* < 0.0001; t-test).

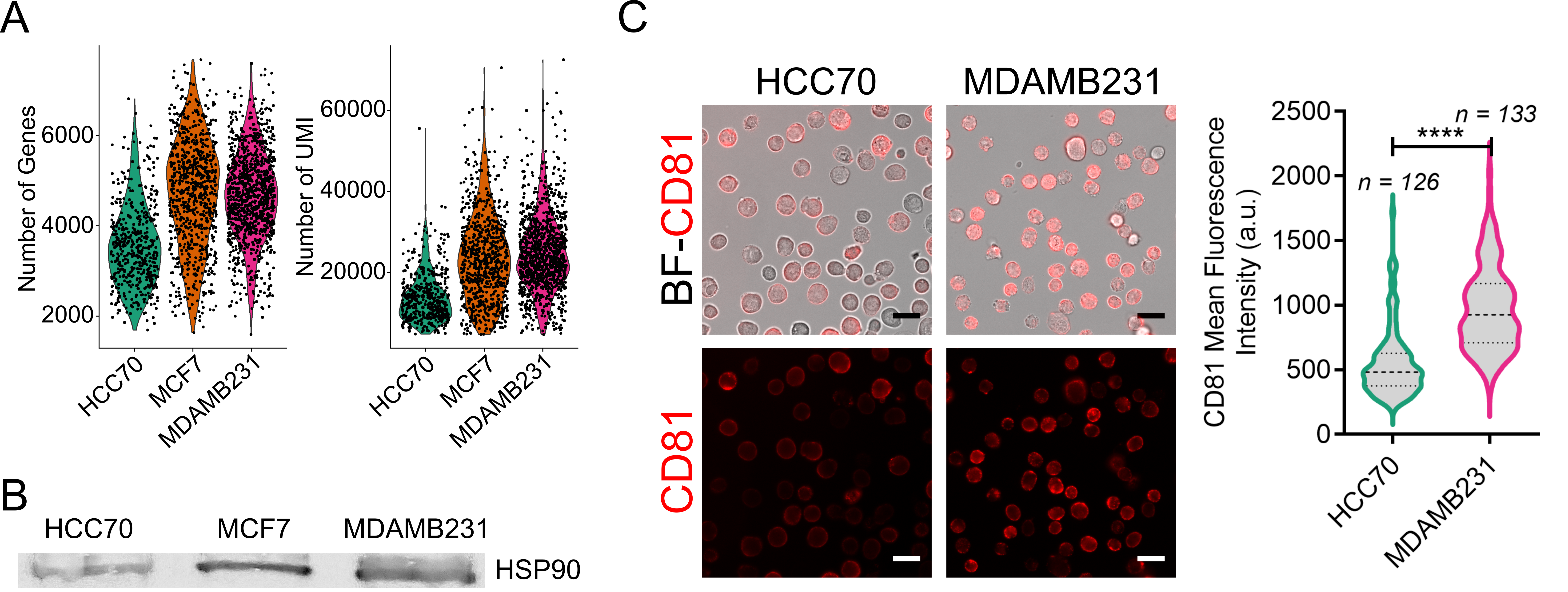

Figure S3. The scRNA-seq analysis and protein expression of MDAMB231, MCF7, and HCC70 cell lines

A. Violin plot of the number of genes and transcriptome per cells.
B. Western blot of HSP90 for HCC70, MCF7 and MDAMB231 cell lines.
C. Left: CD81 surface expression of HCC70 and MDAMB231 using fluorescence microscopy. Scale bar is 25 µm. Right: Violin plots of the median and quantiles of CD81 intensity (**** *p* < 0.0001; t-test).

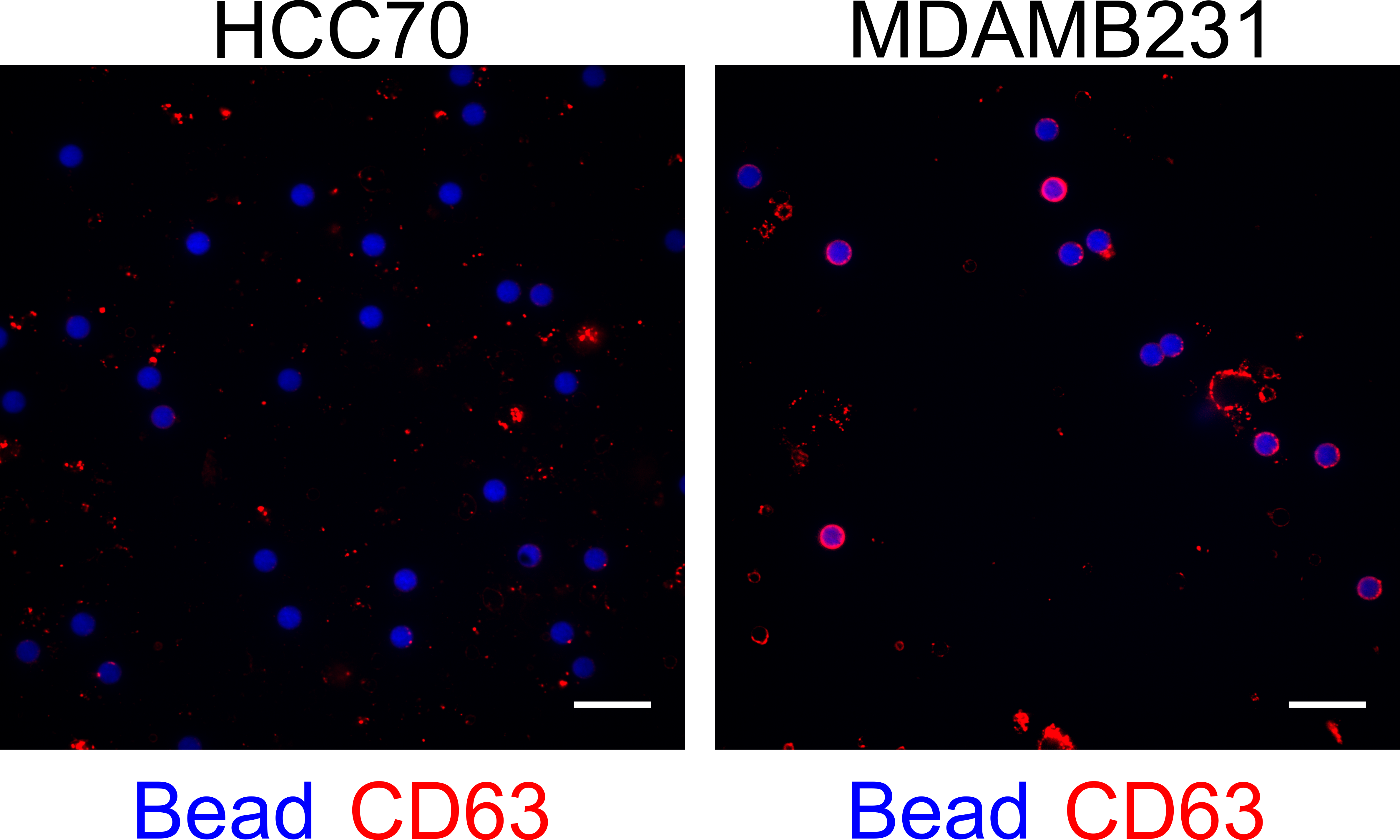

Figure S4. Overlay images of EVs captured onto beads upon co-culture with the appropriate cell lines using transwell assay.

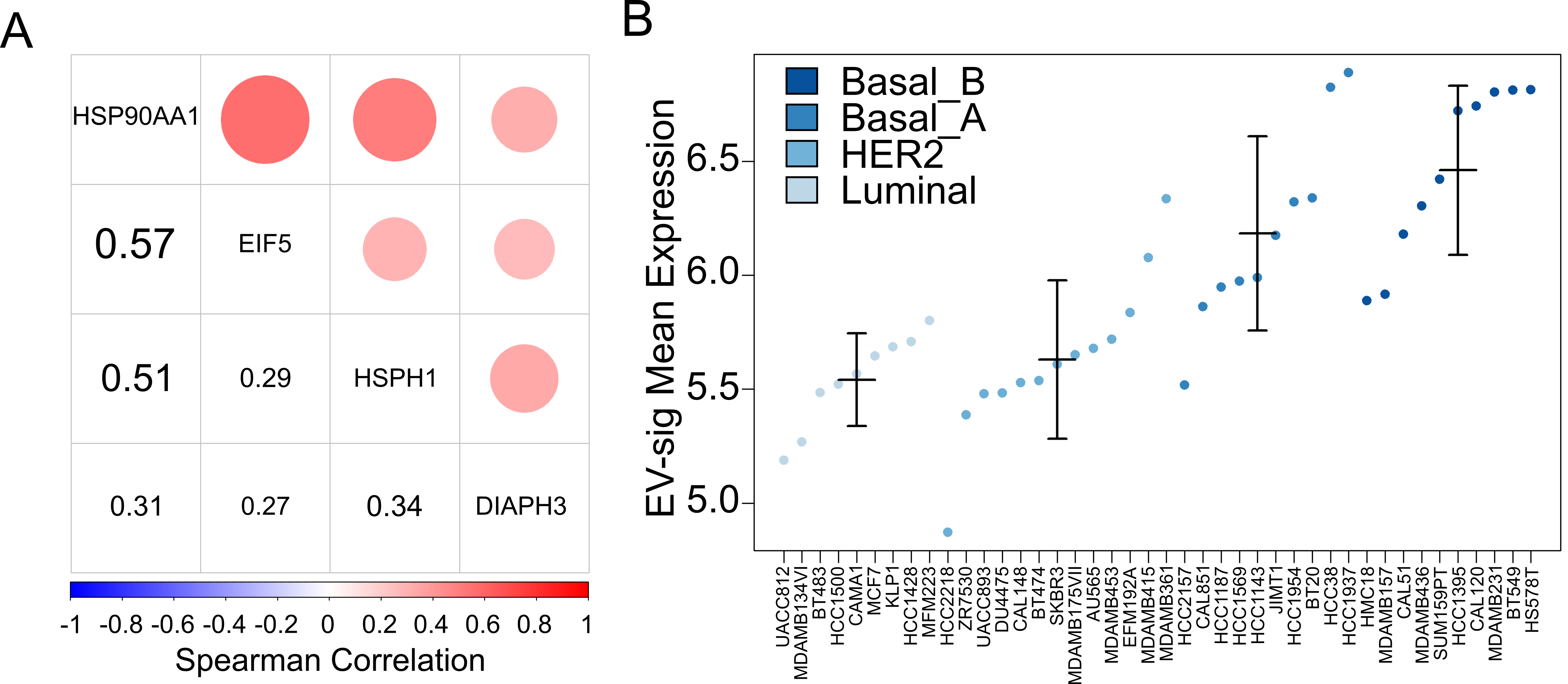

Figure S5. The prediction of EV secretion in cell lines of the CCLE dataset.

A. Spearman correlation coefficient for four core EV genes in 1304 cell lines available on the CCLE dataset.
B. Average expression of the top genes identified from MDAMB231-S cells in breast cancer cell lines available on the CCLE dataset. The cell lines are grouped based on four breast cancer subtypes, luminal, HER2-enriched, basal A, and basal B.

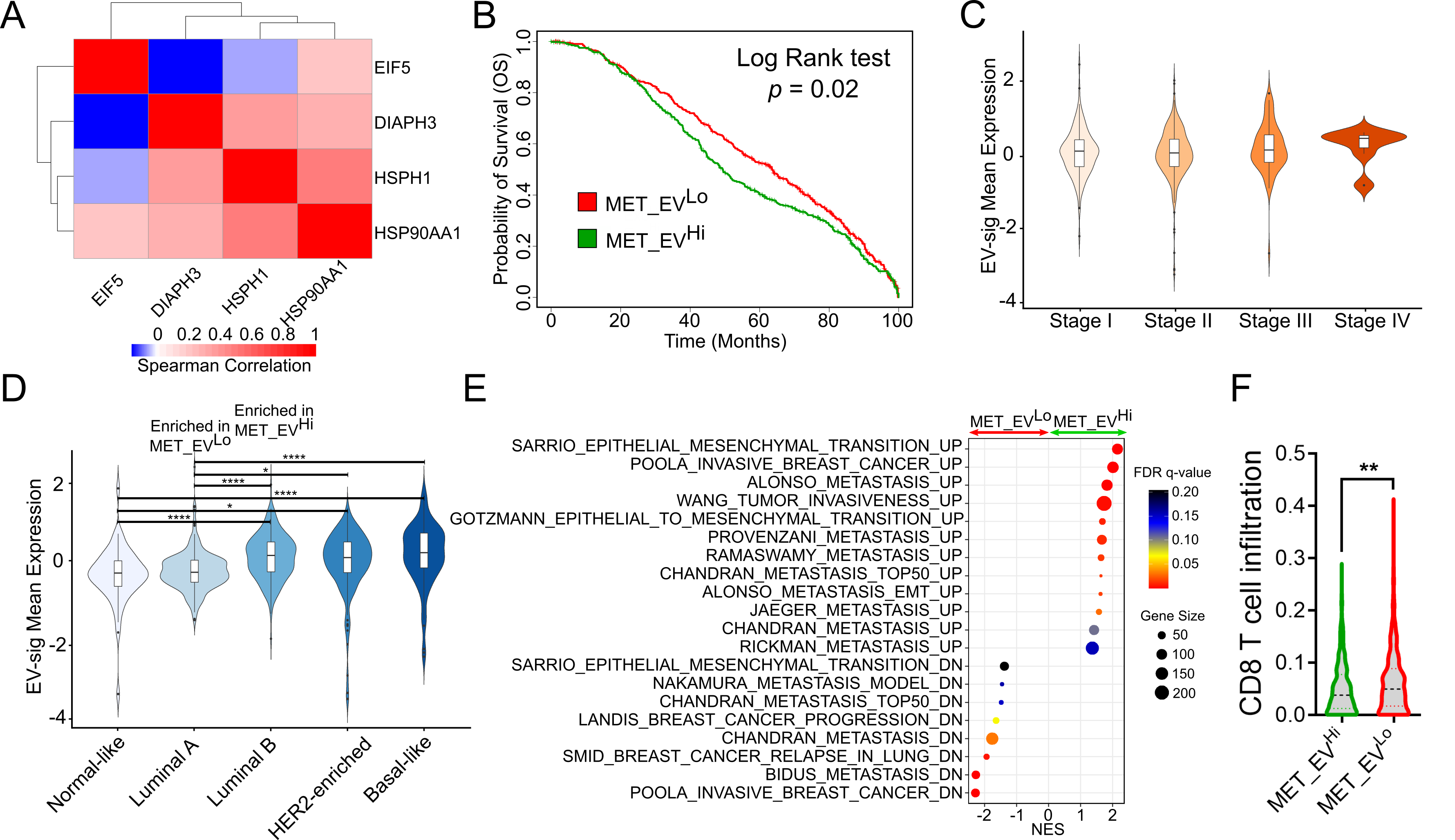

Figure S6. The expression of EV-sig genes is correlated with poor survival in METABRIC dataset.

A. Spearman correlation coefficient for four core EV genes in breast cancer patients.
B. Overall survival of breast cancer patients divided by the median of the average expression of EV-sig genes (n = 405, and 404 for MET_EV^Hi^ and MET_EV^Lo^, respectively)
C. Violin plots of average expression of EV-sig genes by breast cancer stage (n=133, 373, 75, and 7 for Stage I to V, respectively; * *p* < 0.05, and **** *p* < 0.0001; one-way ANOVA).
D. Violin plots of average expression of EV-sig genes by the breast cancer subtype (n=55, 230, 216, 118, and 103 for normal-like, luminal A, B, HER2-enriched, and basal-like subtypes, respectively; * *p* < 0.05, and **** *p* < 0.0001; one-way ANOVA).
E. Normalized enrichment scores (NESs) of pathways associated with metastasis in patients with high and low levels of EV-sig gene expression.
F. CD8^+^ T cells infiltration based on CIBERSORTx data for patients with high and low levels of EV-sig gene expression (** *p* < 0.01; t-tests).

**Table S1: DEGs upregulated in MDAMB231-S compared to MDAMB231-NS (provided in a separate file)**

**Table S2: List of DEGs upregulated in MDAMB231-S compared to MDAMB231-NS with/without overlap** **with mRNA and proteins from the ExoCarta dataset (provided in a separate file)**

**Table S3: The link between top 34 genes and cancer EV**

|  | Linked to Cancer | Linked to Breast Cancer | References |
| --- | --- | --- | --- |
| FTL | Y | Y | [1] |
| TIMP3 | Y | N | [2] |
| SNAPC1 | N | N | - |
| FXYD5 | N | N | - |
| G0S2 | N | N | - |
| ACTN1 | Y | N | [3] |
| RPL21 | Y | N | [4] |
| LCP1 | Y | Y | [5] |
| HMGB1 | Y | N | [6] |
| POMP | N | N | - |
| OGFRL1 | N | N | - |
| HSP90AA1 | Y | Y | [7] |
| GTF3A | Y | N | [2] |
| TPP2 | Y | N | [6] |
| NQO1 | Y | N | [8] |
| IPO5 | Y | N | [9] |
| ADM | Y | N | [10] |
| STXBP6 | Y | N | [6] |
| UCHL3 | Y | N | [11] |
| TFDP1 | N | N | - |
| TPT1 | Y | N | [2] |
| HSPH1 | Y | N | [3] |
| EIF5 | Y | N | [6] |
| FTH1 | Y | Y | [1] |
| SAP18 | Y | N | [6] |
| UBL3 | Y | Y | [12] |
| ZFP36L1 | N | N | - |
| DBN1 | Y | N | [13] |
| CAV1 | Y | Y | [14] |
| SKA3 | N | N | - |
| DIAPH3 | Y | N | [11] |
| DNAJC15 | N | N | - |
| ACTG1 | Y | Y | [15] |
| MRPL57 | N | N | - |

**Table S4: DEGs upregulated in MDAMB231, MCF7 and HCC70 cell lines (provided in a separate file)**
